# The WOPR family protein Ryp1 is a key regulator of gene expression, development, and virulence in the thermally dimorphic fungal pathogen *Coccidioides posadasii*

**DOI:** 10.1101/2021.07.26.453804

**Authors:** M. Alejandra Mandel, Sinem Beyhan, Mark Voorhies, Lisa F. Shubitz, John N. Galgiani, Marc J. Orbach, Anita Sil

**Affiliations:** School of Plant Sciences, University of Arizona, Tucson, AZ 85721, USA; Valley Fever Center for Excellence, The University of Arizona, Tucson, AZ 85724, USA; Department of Microbiology and Immunology, University of California San Francisco, San Francisco, CA 94143, USA; Department of Infectious Diseases, J. Craig Venter Institute, La Jolla, CA 92037, USA; Department of Medicine, University of California San Diego, La Jolla, CA 92037, USA

**Keywords:** *Coccidioides posadasii*, morphology, virulence, dimorphic fungi

## Abstract

*Coccidioides* spp. are mammalian fungal pathogens endemic to the southwestern US and other desert regions of Mexico, central and South America, with the bulk of US infections occurring in California and Arizona. In the soil, *Coccidioides* grows in a hyphal form that differentiates into 3-5 micron asexual spores (arthroconidia). When arthroconidia are inhaled by mammals they undergo a unique developmental transition from polar hyphal growth to isotropic expansion with multiple rounds of nuclear division, prior to segmentation, forming large spherules filled with endospores. Very little is understood about the molecular basis of spherule formation. Here we characterize the role of the conserved transcription factor Ryp1 in *Coccidioides* development. We show that *Coccidioides* Δ*ryp1* mutants have altered colony morphology under hypha-promoting conditions and are unable to form mature spherules under spherule-promoting conditions. We analyze the transcriptional profile of wild-type and Δ*ryp1* mutant cells under hypha- and spherule-promoting conditions, thereby defining a set of hypha- or spherule-enriched transcripts (“morphology-regulated” genes) that are dependent on Ryp1 for their expression. Forty percent of morphology-regulated expression is Ryp1-dependent, indicating that Ryp1 plays a dual role in both hyphal and spherule development. Ryp1-dependent transcripts include key virulence factors such as SOWgp, which encodes the spherule outer wall glycoprotein. Concordant with its role in spherule development, we find that the Δ*ryp1* mutant is completely avirulent in the mouse model of coccidioidomycosis, indicating that Ryp1-dependent pathways are essential for the ability of *Coccidioides* to cause disease. Vaccination of C57BL/6 mice with live Δ*ryp1* spores does not provide any protection from lethal *C. posadasii* intranasal infection, consistent with our findings that the Δ*ryp1* mutant fails to make mature spherules and likely does not express key antigens required for effective vaccination. Taken together, this work identifies the first transcription factor that drives mature spherulation and virulence in *Coccidioides*.

**Author Summary:** *Coccidioides* species, *C. immitis* and *C. posadasii*, are dimorphic fungal pathogens that commonly infect humans in North, Central, and South America, causing the respiratory fungal disease known as Valley Fever. *Coccidioides* grows as hyphae in the soil and differentiates into unique structures called spherules in the mammalian host. Spherules expand and internally divide to form endospores, which are released to facilitate dissemination of the pathogen within the host. The mechanisms underlying spherule differentiation remain largely unknown. In this study, we generated knockout mutants (Δ*ryp1*) of the conserved transcription factor Ryp1 in *C. posadasii* and characterized its role in spherule formation and virulence. We found that Ryp1 is required for the formation of mature spherules and colonization of mouse lungs. Transcriptional profiling of the Δ*ryp1* mutant and the wild-type strain shows that Ryp1 regulates the expression of a subset of the transcripts that are either upregulated in wild-type spherules or in wild-type hyphae. These findings suggest that Ryp1 has a dual role in regulating morphology and virulence under host conditions as well as regulating genes involved in hyphal growth in the environment.

## Introduction

*Coccidioides spp*. are fungal pathogens endemic to California, Arizona, and other desert regions in the Americas [1]. *Coccidioides* infects otherwise healthy individuals when they inhale spores from the soil. The prevalence of *Coccidioides* infection rose 9-fold between 1998 and 2011 and has continued to spike, with incidence now at an all-time high [2]. Elegant observational studies of *Coccidioides* have established its complex life cycle, its disease etiology, and its interaction with the mammalian immune system. The molecular basis of these attributes remains poorly understood, although candidate-based studies have implicated a handful of *Coccidioides* genes in growth and virulence.

*Coccidioides* is one of a group of thermally dimorphic fungal pathogens that grow in a sporulating hyphal form in the soil [3]. Upon introduction into the mammalian host, spores undergo differentiation into a parasitic form. Indeed, the defining characteristic of *Coccidioides* pathogenesis is development of the spore (arthroconidium) into a multicellular structure called the spherule [1,4]. In spherule development, instead of germinating and growing as polar hyphae, the arthroconidium undergoes isotropic enlargement in the mammalian lung with multiple rounds of mitosis prior to formation of internal spores (endospores) resulting in a 60 to 100 μm micron multi-cellular spherule, encased by a spherule outer wall [5]. If a spherule ruptures, endospores are released and can re-initiate the spherule cycle, either locally or following dissemination to other sites within the host. Known virulence genes in *Coccidioides* are expressed concomitant with spherule development [6–9]. One critical unanswered question in *Coccidioides* biology is the nature of the regulatory network that drives spherulation and virulence.

To identify putative regulators of spherulation, we assessed the role of the conserved fungal transcription factor, Ryp1, that is involved in development of the parasitic form in response to temperature for the other thermally dimorphic fungi. We had previously shown that the WOPR transcription factor Ryp1 is a master regulator that is required for the formation of the host form in the thermally dimorphic fungal pathogen *Histoplasma* [10,11]. Additionally, Ryp1 is a master regulator of gene expression in response to temperature in *Histoplasma*, as it is required for the vast majority of the gene expression program at 37°C [10,11]. Investigating the role of Ryp1 in *Coccidioides* was particularly compelling since Ryp1 orthologs are found throughout the fungal kingdom and are required for key developmental transitions in numerous fungi [12–16]. For example, the *Candida albicans* ortholog Wor1 regulates cell-type specification by driving the switch from “white” cells to “opaque” cells [17,18].

Here we deleted the *RYP1* gene in *Coccidioides posadasii* and showed that the resultant mutant is unable to undergo mature spherulation. We used RNA-seq to show that Ryp1 has a role in gene expression in both spherules and hyphae. The Δ*ryp1* mutant failed to express the normal complement of hypha-enriched transcripts, and most notably, the immature Δ*ryp1* spherules expressed only a subset of normal spherule-enriched genes, consistent with their inability to fully differentiate. The Δ*ryp1* mutant was completely avirulent in the mouse model of *Coccidioides* infection, demonstrating that the Ryp1 transcription factor is a key regulator that links the ability to form mature spherules and the expression of critical virulence traits.

## Results

### Deletion of *Coccidioides posadasii RYP1*

To define potential regulators of *Coccidioides* parasitic phase development, we identified the *C. posadasii* ortholog of the conserved regulator of fungal development, *RYP1*, based on similarity to the *H. capsulatum* Ryp1 protein [10]. Ryp1 is a member of the WOPR family of transcription factors. The *C. posadasii* strain Silveira *RYP1* gene, CPSG_00528, encodes a 404 amino acid protein that is 68% similar to the 487 *H. capsulatum RYP1* protein [10], with 84% identity and 91% similarity over the N-terminal half of the protein including the WOPRa and WOPRb regions and the putative nuclear localization signal). To determine the role of *RYP1* in the *C. posadasii* life cycle, gene replacement strains were made via *Agrobacterium* transformation [19,20]. Transformed lines were screened for homologous recombination of a deletion construct where the *E. coli hphB* gene was cloned between 1.2 kb 5’ and 3’ flanking regions of the *RYP1* gene, using an approach similar to that used for creation of the Δ*cps1* strain [19]. A total of 40 transformed lines were generated between two transformations with 42% being *RYP1* gene replacements as analyzed by DNA hybridization for the first set of transformants and PCR for the second set (data not shown). The *RYP1* gene replacement strains (Δ*ryp1*) have a single insertion of the 1.4 kb *E. coli hphB* expression cassette replacing the full 1.3 kb *RYP1* coding region. Phenotypic analyses of Δ*ryp1* mutants were performed using several independent transformed strains and compared to the parental Silveira strain (WT) and one strain (1563.19) where the *RYP1* deletion construct had integrated randomly in the genome leaving the *RYP1* gene intact.

### Ryp1 is required for *Coccidioides* development

Deletion of *RYP1* resulted in several *in vitro* phenotypes (Fig 1). The Δ*ryp1* arthroconidia were delayed in germination, with visible colonies appearing after 7 days at 25°C on 2X GYE media while the progenitor WT and the ectopic transformed strains gave visible colonies three days after plating. The radial growth of Δ*ryp1* colonies was also retarded relative to WT or ectopic transformed strains (Fig 1G) and Δ*ryp1* colonies failed to grow to the edge of petri dishes (Fig 1A-D). The Δ*ryp1* strains exhibited a granular phenotype at the edges of the colony (Fig 1E and F), possibly indicative of more dense conidiation. The most dramatic phenotype was observed under *in vitro* spherulation conditions: the Δ*ryp1* mutant failed to differentiate into mature spherules. Instead the arthroconidia remained as thin-walled structures at all time points and by 72 hours had formed short hyphal-like segmented structures (Fig 2).

**Figure 1.**
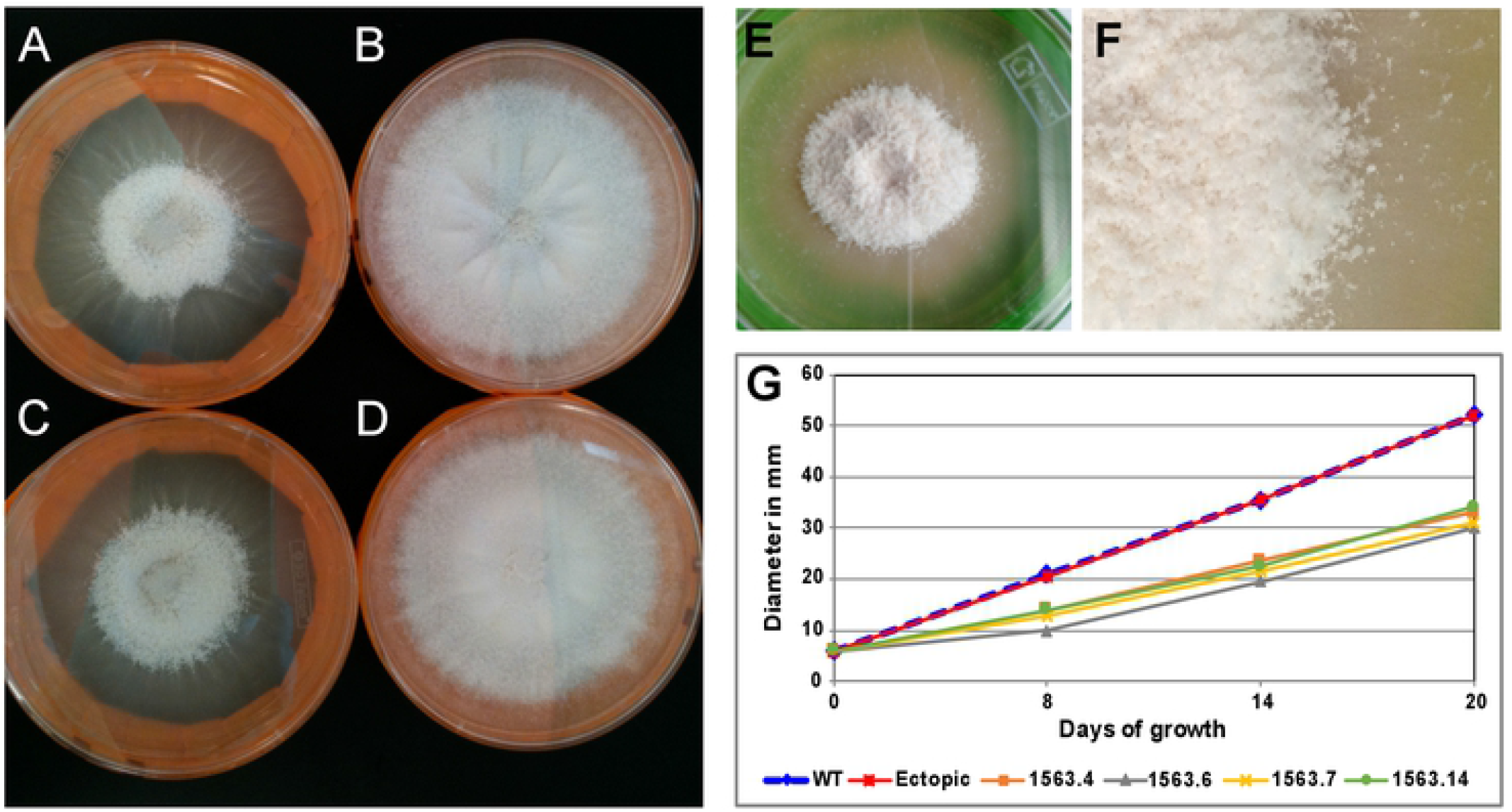
Growth morphology of Δ*ryp1* mutants. *C. posadasii* Δ*ryp1* strains exhibit a growth-limiting phenotype (A, C), in comparison to WT (B) and an ectopic transformant (D). (E) The Δ*ryp1* strains have a granular phenotype. (F) Enlarged view of the Δ*ryp1* colony edge is shown. (A-F) All colonies were inoculated at the center of the agar media and grown at room temperature for 28 days. Representative image of each strain is shown. (G) Plugs of 6 mm were transferred from freshly growing plates to 2X GYE plates and incubated at room temperature, to measure colony radial growth. Colony diameters were measured on days 8, 14 and 20. Four independent Δ*ryp1* strains (1563.4, 1563.6, 1563.7 and 1563.14), as well as the WT parental strain and an ectopic transformed strain (1563.1) were tested.

**Figure 2:**
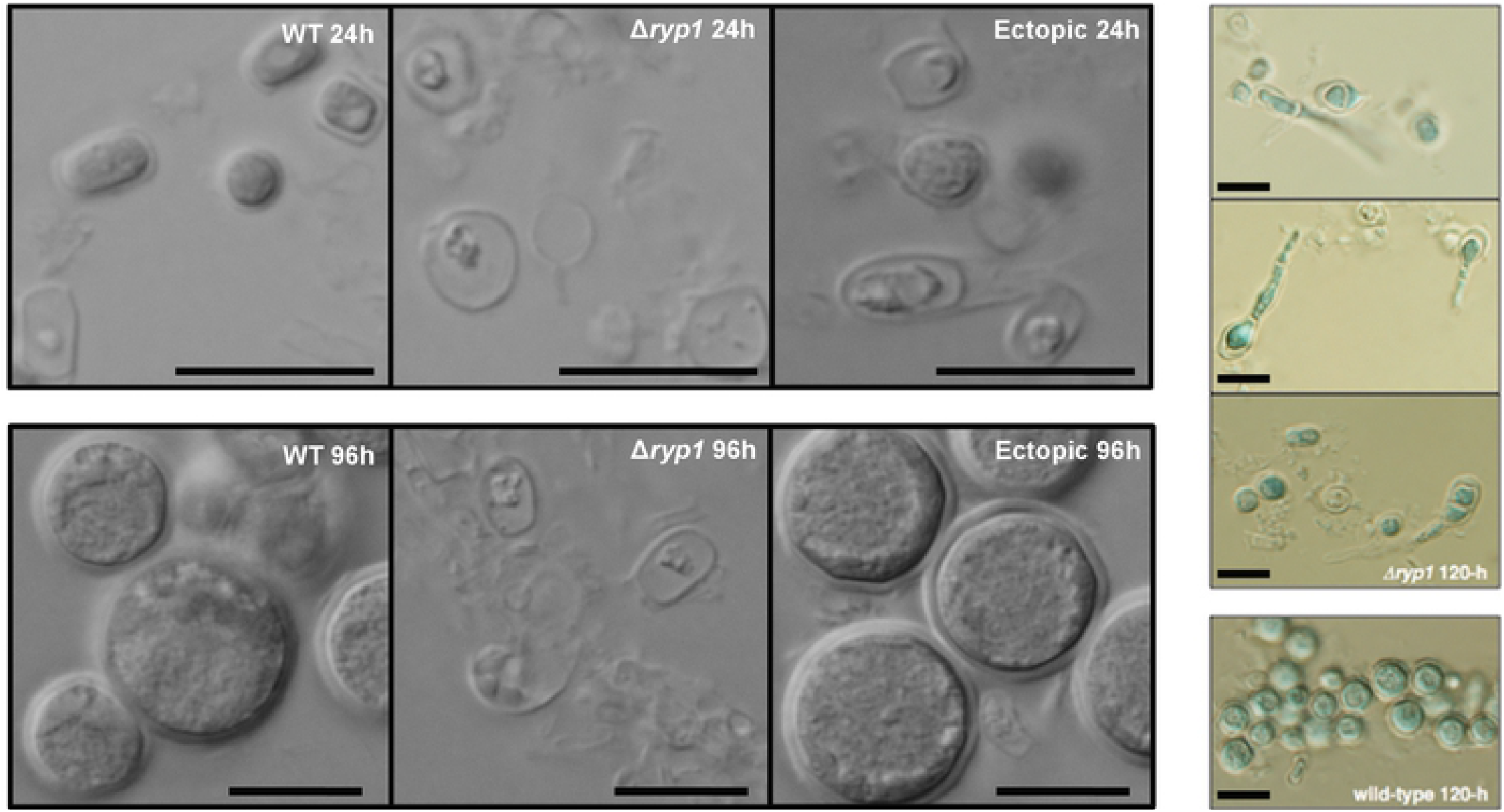
*C. posadasii* Δ*ryp1* does not produce normal spherules *in vitro*. Spherules of WT, a Δ*ryp1* mutant and an ectopic transformed strain, grown in Converse medium at 37°C with 20% CO_2_ and aliquots analyzed during *in vitro* spherulation. Images at 24 h show that while the WT and ectopic strains are rounding up to form early spherules, Δ*ryp1* does not form round structures. At 96 h and 120 h, the WT and ectopic strains produced mature spherules, while the Δ*ryp1* exhibited limited polar hyphal growth. Black bars represent 10 µm.

### Ryp1 regulates distinct sets of genes in spherules and hyphae

Our previous studies revealed that *H. capsulatum* Ryp1 is required for yeast-phase growth and is responsible for the expression of the majority of yeast-phase specific genes *[10]*. Given that *C. posadasii* Δ*ryp1* mutants are defective in spherulation, we postulated that *C. posadasii* Ryp1 is also responsible for transcriptional regulation of morphology-associated genes. To this end, we performed RNA-seq to transcriptionally profile WT and Δ*ryp1* strains of *C. posadasii* grown under either spherulation or hyphal conditions. Under our experimental conditions, 9711 transcripts were observable (transcripts per million, TPM, ≥ 10) in at least one sample. Of these, 4175 transcripts (about 43% of all detected transcripts) were significantly differentially expressed (Fig 3) with at least a two-fold change in at least one of the following three comparisons: (1) a comparison of WT spherules and WT hyphae (WT^Sph^/WT^Hy^) identified 2985 transcripts that we refer to as “morphology-regulated” genes; (2) a comparison of WT vs Δ*ryp1* under spherule conditions identified 1742 Ryp1-dependent transcripts; and (3) a comparison of WT vs. Δ*ryp1* under hyphal conditions identified 925 Ryp1-dependent transcripts (Fig 3B). Not all of these Ryp-1 dependent transcripts in comparisons 2 and 3 are morphology-regulated genes, and in fact, 1157 morphology-regulated transcripts (39% of the 2985 WT^Sph^/WT^Hy^ comparison) were dependent on Ryp1 either under spherulation or hyphal conditions. Thus, in contrast to *H. capsulatum*, over half of the morphology-regulated expression is Ryp1-independent, and Ryp1 regulates distinct sets of genes whose expression correlates with either spherule or hyphal growth conditions. To further explore the 4175 differentially expressed transcripts, we grouped them in 17 classes by expression pattern (Fig 3A, S1 Table). Ryp1-dependent transcription under spherulation conditions is correlated with WT morphology-regulated transcription: spherule-enriched genes are more likely to be Ryp1-induced (class 2 in Fig 3A) while hypha-enriched genes are more likely to be Ryp1-repressed in spherules (Fig 3C and class 4 in Fig 3A). In contrast, Ryp1-dependent transcription under hyphal conditions is not correlated with morphology-regulated transcription, as can be seen from the shape of the scatter plot (Fig 3D).

**Figure 3.**
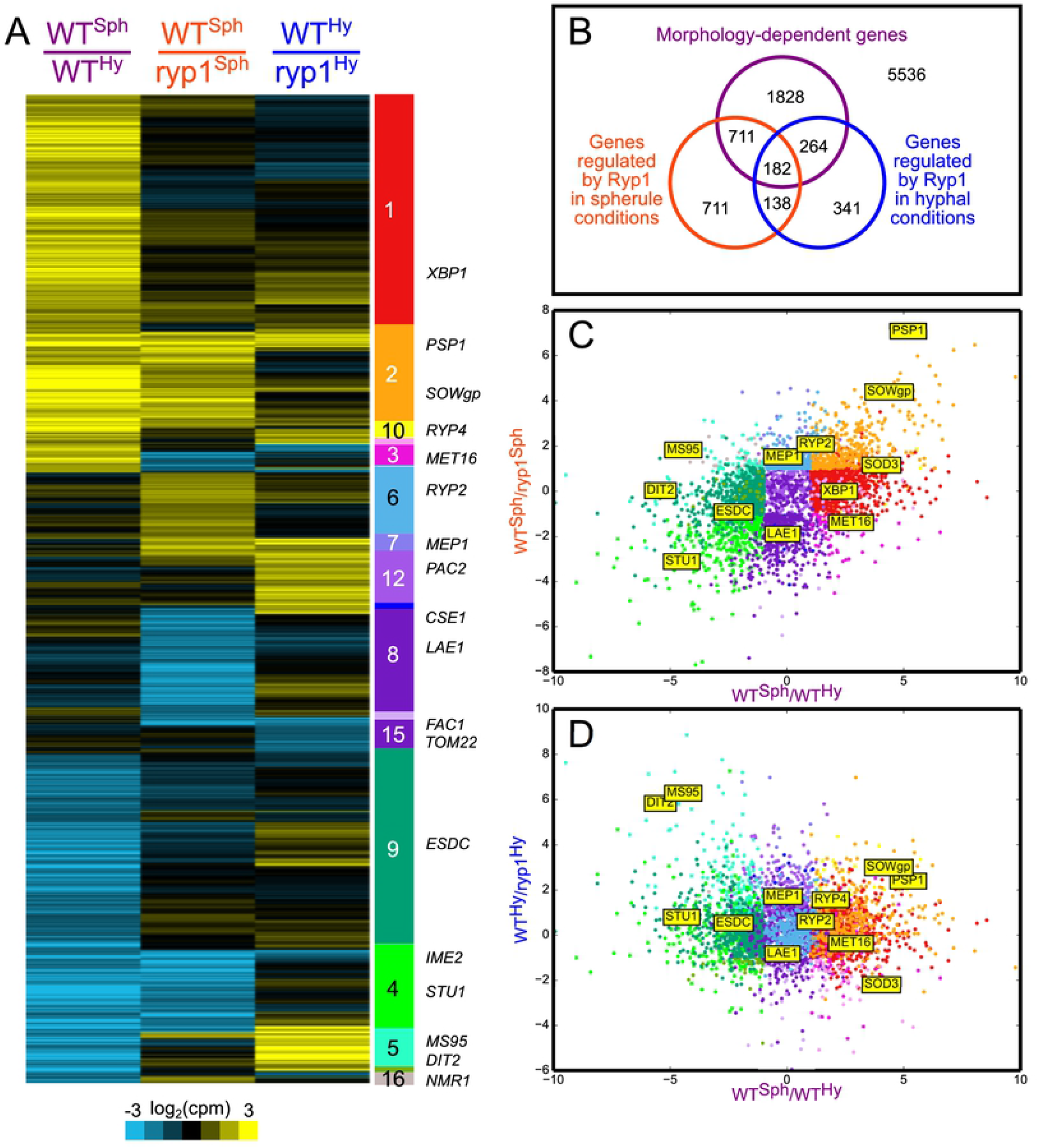
Ryp1 regulates distinct sets of genes in spherules and hyphae. (A) Heatmap showing differential expression of the 4175 genes described in the text. Genes are grouped by expression pattern (significantly up, down, or neutral in each of the three contrasts) with different classes labeled in the color bar to the right of the heatmap. The position of genes of interest in the heatmap are indicated to the right of the color bar. (B) Venn diagram of genes observed to be morphology-regulated (differentially expressed in WT^Sph^/WT^Hy^), Ryp1-dependent in spherulation conditions (differentially expressed in WT^Sph^/Δ*ryp1*^Sph^), or Ryp1-dependent in hyphal conditions (differentially expressed in WT^Hy^/Δ*ryp1*^Hy^). (C and D) Scatter plots of the differentially expressed genes comparing the WT spherule-enrichment (right) or hyphal-enrichment (left) to *ryp1*-depletion (top) or *ryp1*-enrichment (bottom) in (C) spherules or (D) hyphae. Genes in the scatter plots are colored by expression pattern, as in the color bar of (A).

### Ryp1 is required for the expression of a set of spherule-enriched transcripts

Early subtractive cDNA hybridization experiments identified four spherule-specific genes (*ALD1, OPS1, PSP1*, and *MDR1*) [21]. Of these, we found that *ALD1* is spherule-enriched and Ryp1-independent (S1 Table, class 1) while the remaining three are both spherule-enriched and Ryp1-dependent at 37°C (S1 Table, class 2). Altogether, 406 (25%) of the 1593 transcripts that are more highly expressed in wild-type spherules versus hyphae are dependent on Ryp1 for their expression. This set includes SOWgp, the best characterized *Coccidioides* virulence factor [22] and CPSG_05265 (CIMG_00509), a previously noted spherule enriched gene [6] at the boundary of a region of introgression between *C. immitis* and *C. posadasii* [23].

Intriguingly, about 5% (84 of 1593) of the spherule-enriched transcripts are more highly expressed in the Δ*ryp1* mutant than WT under spherulation conditions, (S1 Table, class 3), indicating their expression is normally repressed by Ryp1 in wild-type spherules. This set includes the sulfur assimilation pathway (*MET3, MET14, MET16*, and *MET10*) required to reduce intracellular sulfate to sulfide. In contrast, we found that the downstream genes *MET5* (the beta subunit to *MET10*) and *MET17A* (which catalyzes the first step of cysteine and methionine biosynthesis) are both spherule-enriched but Ryp1-independent, while the genes acting in opposition to sulfur assimilation (both the sulfite oxidase *SOX1* and the sulfite exporter *SSU1*) are spherule-enriched and Ryp1-dependent. These data are consistent with Ryp1 regulation promoting sulfur efflux versus influx.

### Ryp1 regulates the expression of a set of hypha-enriched transcripts

Analysis of the hypha-enriched transcripts showed that about 25% of this set (351 out of 1392) are Ryp1-repressed when cells are grown under spherulation conditions (S1 Table, class 4). Notably, this set includes the APSES transcription factor *STU1*, which regulates hyphal morphology in *Histoplasma* [24] and has conserved hyphal enrichment in *C. immitis* [8] and all of the thermally dimorphic pezizomycetes for which RNA-seq data is available [25]. Additional genes in this set consistent with hyphal growth include 5 septins (*CDC10, CDC12, CDC3, ASPA*, and *ASPE*), which are cytoskeletal proteins involved in cell cycle regulation [26]. Intriguingly, 33 of the 351 genes in this set are also Ryp1-dependent under hyphal growth conditions. In particular, the *Coccidioides* ortholog of the OefC transcription factor, which confers a “fluffy” phenotype in *Aspergillus nidulans* when overexpressed [27], exhibits this Ryp1-dependent regulation in hyphae. Furthermore, about 12% (164 of the 1392) of the hypha-enriched transcripts are found to be Ryp1-dependent under hyphal-promoting conditions (S1 Table, class 5). This set includes the stress response gene *DDR48*/*MS95*, which is highly enriched in *Histoplasma* hyphae [28], as well as both *DIT2*, an enzyme involved in the production of N,N-bisformyl dityrosine, and the bisformyl dityrosine transporter *DTR1*. In *Saccharomyces cerevisiae*, bisformyl dityrosine is a major building block of the protective ascospore cell wall, whose assembly depends on both *DIT2* [29] and *DTR1* [30]. These Ryp1-dependent hypha-enriched genes also include many enzymes with NADH cofactors, including two (CodA and AifA) with roles in the response to reductive stress [31,32]. Taken together, our results suggest that Ryp1 has a dual role in *Coccidioides* development, regulating the expression of subsets of both spherule- and hypha-enriched genes.

### Ryp1 regulates expression of morphology- or temperature-independent transcripts

In *Histoplasma*, Ryp1 directly interacts with two other transcriptional regulators, Ryp2 and Ryp3, to regulate yeast-phase growth [11]. The orthologs of *RYP2* and *RYP3* in *Coccidioides* are not differentially regulated in wild-type spherules and hyphae; however, *RYP2* displays a Ryp1-dependent expression pattern under spherulation conditions (S1 Table, class 6). In addition, 282 transcripts show a similar morphology-independent but Ryp1-dependent expression pattern under spherulation conditions, including the cytosolic catalase *CATP*, the tyrosinase *TYR2*, as well as ∼40 genes involved in primary carbohydrate, nucleotide, and amino acid metabolism. An additional 75 transcripts display a morphology-independent but Ryp1-dependent expression pattern under both spherule- and hypha-promoting conditions (S1 Table, class 7). Among these genes are *MEP1*, a protease that contributes to immune evasion via degradation of SOWgp [33]; urease (CPSG_08438), a known virulence factor of *Coccidioides [34]*; *UAZ1*, a uricase that catalyzes the first step in the breakdown of uric acid; and *CTR2*, a copper transporter observed to be spherule-enriched in *C. immitis*.

In addition to the Ryp1-dependent transcripts, there are 432 transcripts that are equivalently expressed in WT spherules and hyphae but are upregulated in Δ*ryp1* mutants compared to wild-type under spherulation conditions (S1 Table, class 8), suggesting that Ryp1 represses their expression. This set includes *LAE1*, a master regulator of secondary metabolism [35]; orthologs of the *Aspergillus* developmental regulators NsdD (*NSD4*) and FlbC (*FBC1*); the MAPK pathway components *STE11, PBS2*, and *SDP1*; and the histidine kinase PhkB (*PHK2*). In *Histoplasma, PHKB* shares a Ryp1-associated divergent promoter with the histidine kinase *PHKA*, and both genes are Ryp1-induced in *Histoplasma* yeast [11]. In *Coccidioides*, these genes likewise share a divergent promoter, and have correlated expression profiles, but the differential expression of *PHKA* is too modest to pass our 2-fold change criterion.

### A significant portion of the morphology-regulated genes are expressed independently of Ryp1

In addition to the spherule-enriched genes that were regulated by Ryp1, there was a significant portion of spherule-enriched genes that were independent of Ryp1. In fact, of 1593 transcripts that are upregulated in wild-type spherules compared to hyphae, 62% of them (1002 transcripts) show Ryp1-independent expression patterns (S1 Table, class 1). Notably, these include genes involved in regulation of morphology: *XBP1*, an APSES transcription factor enriched in *Histoplasma* yeast [24], *SSK1* and *SKN7*, response regulators required for *Histoplasma* yeast morphology (Beyhan et al, unpublished), the redox related genes *TSA1* and *NIR1*, and *HPD1*, which has previously been noted as a morphology related gene in *Coccidioides* [6,9] and *Paracoccidioides* [36]. Similarly, among 1392 hypha-enriched transcripts, 62% (862 transcripts) are Ryp1-independent in our dataset (S1 Table, class 9). These include five enzymes of gluconeogenesis (*PCK1, GAPDH (TDH1), TPI1*, and *FBP1*), as well as genes related in glutamate import, amino acid catabolism, peptidase activity, and protein complex disassembly -- all consistent with hyphae utilizing proteins as a primary carbon source. Notably, *PYC2*, the first enzyme of gluconeogenesis, is both enriched in hyphae and Ryp1-repressed in spherules.

### *RYP1* deletion mutants are avirulent

The critical role of Ryp1 in spherule formation and gene regulation suggested that it may be either reduced in virulence, or avirulent, *in vivo*. To test this, we assessed the virulence of the Δ*ryp1* strain using the mouse model of coccidioidomycosis in C57BL/6 mice. Twelve mice were intranasally infected with 50 or 1,000 arthroconidia of Δ*ryp1* strain 1563.7 and disease progression was compared to infection with 50 WT arthroconidia or 4,400 Δ*cps1* spores which were previously shown to be avirulent [19]. Fungal burden in the lungs and spleen as well as survival was monitored for 28 days post-infection. Additionally, two mice from each group were sacrificed after 12 days for histopathological studies. The WT-infected mice all became moribund and were euthanized between day 12 and 19, while all Δ*ryp1* and Δ*cps1*-infected mice survived to day 28. Lung and spleen homogenates did not yield *Coccidioides* colony-forming units (CFUs) from infections with the Δ*ryp1* and Δ*cps1* mice, consistent with an inability of these mutant strains to survive *in vivo*. In a follow-up experiment, four additional Δ*ryp1* strains were screened for virulence along with an ectopic transformed line (Fig 4). As in the first experiment, Δ*ryp1-*infected mice, which received between 525 and 1133 arthroconidia, survived for the full duration of the 28-day study (Fig 4). In contrast, seven of eight mice infected with 52 spores of the ectopically integrated *RYP1* deletion construct or infected with 49 WT spores died by day 17. One mouse in each of these two groups survived for 28 days but had significant pulmonary disease and dissemination and would have ultimately succumbed to the infection. As we observed previously, there was no growth of *Coccidioides* from the lungs or spleens of the Δ*ryp1-*infected mice. The mean lung CFUs of the mice infected with ectopic strain 1563.1 were 2.87 x 10^7^ (range 1.24 x 10^5^-4.7 x 10^7^), and for those infected with WT were 1.31 x 10^7^ (range 1.06 x 10^5^ – 4.8 x 10^7^), with dissemination to the spleens in all mice as reflected by spleen CFUs.

**Figure 4:**
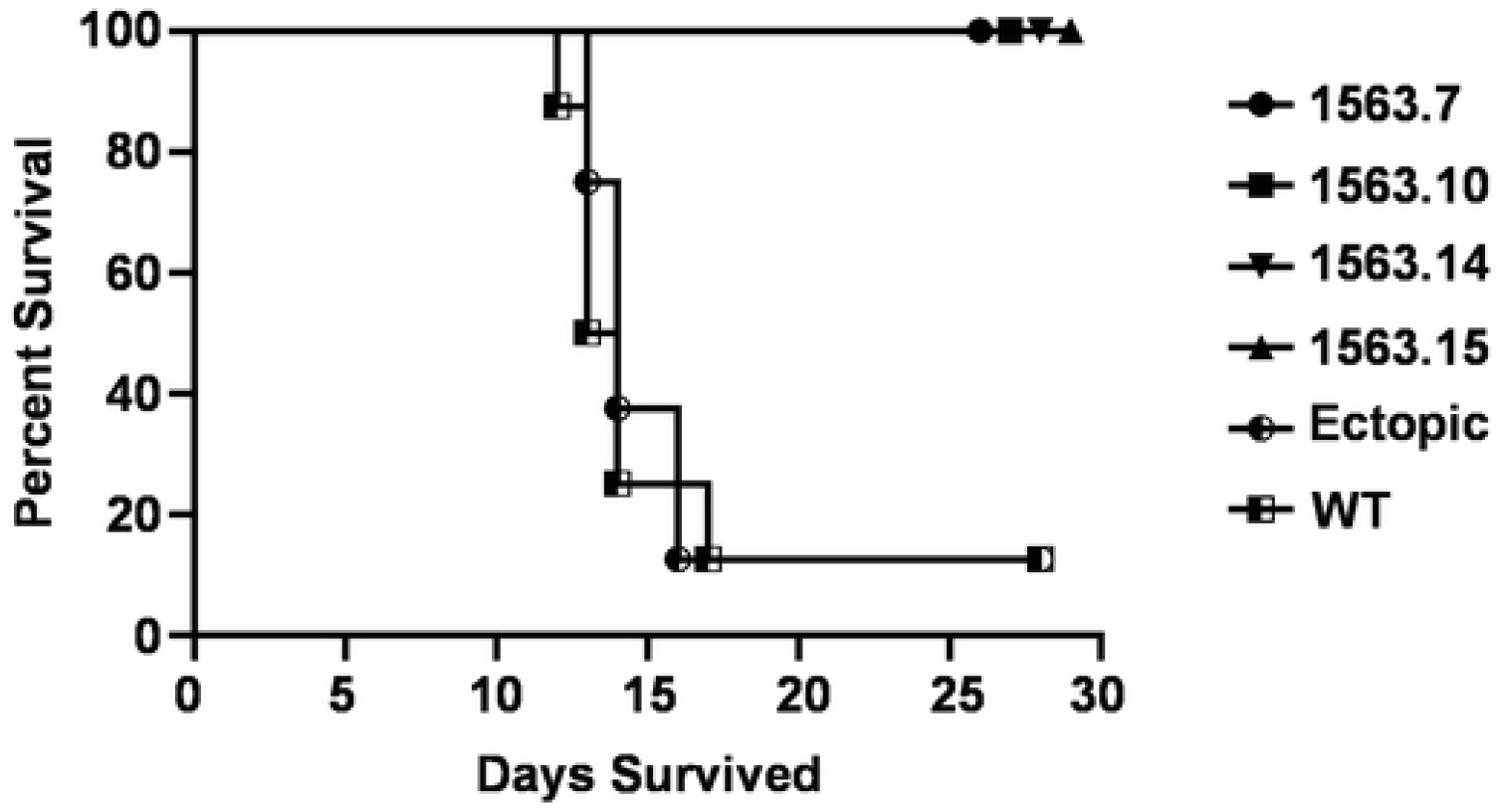
Infection of mice with *C. posadasii* Δ*ryp1*. Survival results for C57BL/6 mice (N=8 mice/group) infected intranasally with four independent Δ*ryp1* strains, 1563.7 (circle), 1563.10 (square), 1563.14 (inverted triangle) and 1563.15 (triangle), one ectopic *ryp1*-transformed strain, 1563.1 (half-filled circle) and WT (half-filled square). The Δ*ryp1* strains were infected at between 525 and 1133 arthroconidia, while 50 arthroconidia of the ectopic stain and WT were used.

### Δ*ryp1* mutants do not provide protection against WT infection

Due to the failure of Δ*ryp1* to cause disease or persist in the lungs of C57BL/6 mice, the Δ*ryp1* strain was tested to determine whether it provides protective immunity against subsequent WT infection, as is the case for the Δ*cps1* strain. The Δ*cps1* vaccine strain and the Δ*ryp1* strain 1563.7 were used to vaccinate C57BL/6 mice either intraperitoneally (IP) or subcutaneously (SC) as described previously [19]. Mice were vaccinated, boosted after two weeks, challenged four weeks later with 90 WT arthroconidia, and sacrificed two weeks later. While Δ*cps1* provided dramatic protection by either SC or IP vaccination, resulting in a 5-log lower lung fungal burden than mice vaccinated with a control adjuvant, vaccination with Δ*ryp1* resulted in no reduction in lung fungal burden and thus, no protection (Fig 5). Mice in the control group had a mean lung fungal burden of 5.3 x 10^6^ CFU, which was similar to that of the Δ*ryp1* SC-vaccinated mice (P<0.05) which had mean lung fungal burdens of 4.1 x 10^6^ CFU and the Δ*ryp1* IP-vaccinated mice with an average of 6.4 x 10^6^ CFU per lung. In contrast, the Δ*cps1*-vaccinated mice had mean lung fungal burdens of 341 CFU for the IP-vaccinated group and 84 CFU for the SC-vaccinated group. *-*Furthermore, when whole spleens were plated for fungal growth, those from Δ*cps1-v-*accinated mice were almost fully free of fungi, with only one spleen of a single Δ*cps1* IP-vaccinated mouse having a small area of growth, likely from a single spherule, appearing seven days after plating, while the control mice and Δ*ryp1-*-vaccinated mice all grew *Coccidioides* by three days after plating. Thus it is clear that the Δ*ryp1* strain does not provide protective immunity against *Coccidioides* infection.

**Figure 5:**
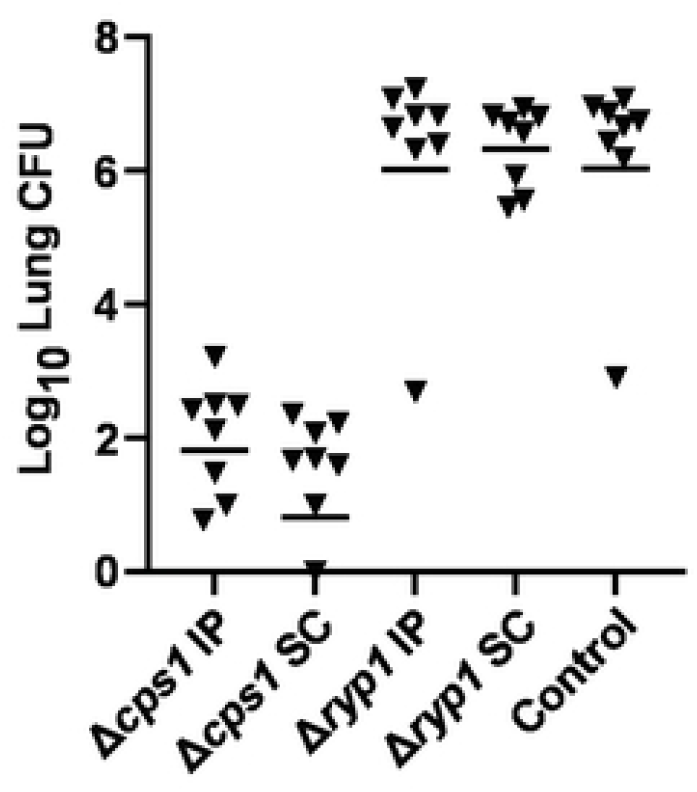
Vaccination of mice with *C. posadasii* Δ*ryp1* mutant. Protection results for C57BL/6 mice vaccinated with either Δ*cps1*, Δ*ryp1* or a *S. cerevisiae* culture supernatant (Control). C57BL/6 mice (N=8 mice/group) were injected either by intraperitoneal (IP) or subcutaneous (SC) injection twice 2 weeks apart with Δ*cps1*, Δ*ryp1*, or Control. Vaccinated mice were challenged with a lethal dose of WT (90 spores) and total lung CFUs were determined 14 days post-infection. Δ*cps1* significantly protects mice from *C. posadasii* infection compared to Control (P<0.004 both comparisons) while Δ*ryp1* shows no protection (P=0.99 vs. control). Bar equals geometric mean.

## Discussion

Spherulation is a critical step of pathogenesis in *Coccidioides* infection, allowing the fungus to adapt to growth within the host and producing endospores which allow expansion and dissemination of the fungus. Here we identified the first transcription factor that is essential for the process of *Coccidioides* spherulation by interrogating the function of Ryp1, a conserved transcriptional regulator that plays a critical role in fungal development and virulence in many fungi, including *Histoplasma*, a close relative of *Coccidioides*. Our results showed that as in *Histoplasma*, Ryp1 is required for development of the host parasitic form; namely spherules in *Coccidioides* and yeast-phase cells in *Histoplasma*. Additionally, in *Coccidioides*, we demonstrated that Ryp1 is absolutely required for virulence in the mouse model of infection, consistent with its critical role in spherulation. The Δ*ryp1* strain failed to differentiate into spherules *in vitro* and failed to persist in susceptible B6 mice, producing no symptoms of disease or viable propagules at 28 days post-infection, even when exposed to 10-20X the level of spores that results in a lethal infection for WT strains. Interestingly, in contrast to *Histoplasma*, where Ryp1 is required for the differential expression of the vast majority of yeast-enriched transcripts, there were a significant number of spherule-enriched genes whose expression was Ryp1-independent, indicating that other transcription factors likely function in parallel to Ryp1 to regulate spherule development. Additionally, *Coccidioides* Ryp1 is required for normal rates of radial growth of hyphal colonies and regulates the expression of a subset of hypha-enriched transcripts. Thus *Coccidioides* Ryp1 has a dual role in both the environmental form and the host form of the fungus where it regulates gene expression and development.

The contrast between *Histoplasma* and *Coccidioides* Ryp1 should be interpreted in the light of the evolutionary relationship between *Histoplasma* and *Coccidioides*--although both are thermally dimorphic fungal pathogens in the order Onygenales, *Histoplasma* is a member of the family Ajellomycetaceae, which also includes the thermally dimorphic pathogens *Paracoccidioides, Blastomyces*, and *Emergomyces*, consistent with a single origin of dimorphism in this family. In contrast, *Coccidioides* is the only known dimorphic member of the family Onygenaceae, and has a unique dimorphic pattern, forming endosporulating spherules during host invasion vs. the more typical hyphal to yeast phase transition. This evolutionary relationship suggests that dimorphism of *Histoplasma* and *Coccidioides* evolved separately, and that some regulatory mechanisms may be shared whereas others are distinct. In *Histoplasma*, since Ryp1 is required for the vast majority of yeast-enriched expression, identifying Ryp-dependent genes by expression analysis of WT vs *ryp1* mutants did not further refine the set of yeast-enriched transcripts defined by comparing the gene expression program of wild-type *Histoplasma* yeast and hyphae [10]. In contrast, only 25% of spherule-enriched transcripts were dependent on *Coccidioides* Ryp1, thereby suggesting that these genes may be key to spherule development and virulence, both of which are defective in the absence of Ryp1. Further identification of key effectors of *Coccidioides* spherulation and virulence could come from future studies identifying direct targets of Ryp1, which was useful in identifying virulence factors in *Histoplasma [11]*.

Transcriptome differences between *Coccidioides* spherules and hyphae have previously been profiled using RNA-seq in both *C. posadasii* and *C. immitis [6,8,37]*. In our dataset, about 30% of the detectable transcripts were differentially regulated between *C. posadasii* spherules and hypha. While there is not a complete overlap with other studies, likely due to differences in growth conditions and stage of spherule development, most of the previously observed highly regulated genes are also differentially regulated in our dataset. For example, SOWgp, the best characterized virulence factor of *Coccidioides* [22], is spherule-enriched in all datasets, and we observe its expression to be Ryp1-dependent. Among the shared hypha-enriched genes in all datasets, *STU1*, an APSES transcription factor that is a regulator of hyphal growth in *Histoplasma [24]*, is Ryp1-repressed in spherules, underscoring the importance of Ryp1 for the expression of morphology-specific genes.

SOWgp, one of the most highly enriched spherule-specific transcripts, is an important component of the *Coccidioides* spherule outer wall (SOW) layer [22]. This lipid-rich layer shed by spherules has been shown to reduce fungicidal activity of neutrophils towards arthroconidia and to promote disseminated disease [38]. The SOW lipids are composed of phospholipids and sphingolipids, with the major sphingolipids being sphingosine and ceramide [38]. In this study, we found that PilA and NCE102, two key factors for assembling eisosomes, punctate membrane associated structures involved in regulating lipid metabolism in response to stress, are enriched in spherules in a Ryp1-dependent manner. Given that eisosomes have a positive role in regulating sphingolipid synthesis [39], and that CPSG_03079, which is orthologous to the ceramide synthase BarA [40], is likewise spherule-enriched and Ryp1-dependent, it is plausible that Ryp1 may regulate production of SOW lipids in spherules.

The transcriptome profiling presented here showed that Ryp1 is responsible for inducing the expression of some spherule-enriched genes and repressing the expression of some hypha-enriched genes. Intriguingly, in addition to these sets of genes, there are other sets of spherule-enriched genes that are not regulated by Ryp1 under spherulation conditions, but are regulated by Ryp1 under hypha-promoting conditions. For example, *RYP4*, which is a major regulator of yeast growth in *Histoplasma*, is only regulated by Ryp1 under hyphal conditions. Here we found that *RYP4* is spherule-enriched in *C. posadasii*, as was also shown in *C. immitis* [8]. However, unlike *Histoplasma*, where *RYP4* expression under yeast-promoting conditions is directly regulated by Ryp1 [11], *RYP4* expression is independent of Ryp1 under spherulation conditions, but dependent on Ryp1 under hyphal conditions (S1 Table, class 10). An additional 67 genes share this expression pattern, including *ACH1*, which is required for detoxification of acetate in *S. cerevisiae* [43] and propionate in *A. fumigatus* [44]. Coexpression of *RYP4* and *ACH1* is notable, given the role of the *RYP4* ortholog FacB in regulation of acetate metabolism in *Aspergillus* [45]. The potential role of Ryp4 in regulating acetate utilization or morphological transitions in *Coccidioides* will be determined by future studies.

In this study, we also explored the role of Ryp1 in pathogenesis using the murine model of coccidioidomycosis. Our results show that Ryp1 is essential for *in vivo* growth and colonization; however, it does not confer protection to the host, unlike the previously characterized Δ*cps1* mutant. Intriguingly, the Δ*cps1* mutant is able to convert to spherules but is impaired in development of mature spherules and endospores [19], whereas the morphology of the Δ*ryp1* mutant *in vivo* is not yet known. Given its morphology defect *in vitro*, we predict that the Δ*ryp1* mutant may not be able to convert to spherules *in vivo*, and thus may not display spherule-specific antigens that are required to elicit a protective host response. Consequently, comparison of the Δ*ryp1* and Δ*cps1* mutants may narrow the search for the molecules responsible for the protective effect of the Δ*cps1* mutant. Furthermore, while the Δ*ryp1* mutant may not be a good vaccine candidate, Ryp1 and other proteins that are required for spherule growth may be effective targets for the development of therapeutics that inhibit the transition from arthroconidia to spherules *in vivo*. Additionally, elucidation of the Ryp1 regulon as described here identifies key downstream effectors that are essential for the development and virulence of the host form of *Coccidioides*, and thus also serve as valuable diagnostic and therapeutic targets.

## Materials and Methods

### Strains and growth conditions

Wild-type *Coccidioides posadasii* (strain Silveira, ATCC 28868) was cultured on 2X GYE medium (2% glucose, 1% yeast extract, and 1.5% agar) at room temperature (approximately 24°C). Mutant strains were selected and maintained on 2X GYE media supplemented with 50 mg/ml of hygromycin, also at room temperature as described previously [19]. Arthroconidia of WT and mutant strains were harvested from 4 week-old cultures, using sterile water by the mini-stir bar method described previously [46] and stored in sterile water at 4°C. Spore numbers were determined with a hemocytometer and viable counts by serial dilution and plating. All manipulations of viable cultures were performed at biosafety level 3 (BSL3). Spherules were prepared by growth in a modified Converse medium at 38°C with 20% CO_2_, with shaking at 180 rpm for 48 hours. To measure colonial radial growth and assess colony morphology, 6 mm plugs of actively growing cultures were transferred to 2X GYE agar plates using transfertubes (Transfertube^®^ Disposable Harvesters, Spectrum^®^ Laboratories,) and grown at room temperature.

### Creation of *RYP1* gene deletion mutants

A gene replacement cassette for the *C. posadasii* strain Silveira *RYP1* gene (CPSG_00528) was constructed using double-joint PCR [47]. DNA fragments of 1.2kb (5’ 1192 and 3’ 1233) representing the 5’ and 3’flanking sequences of CPSG_00528 were amplified by PCR with oligonucleotide primer combinations OAM1153/OAM1154, and OAM1155/OAM1156, respectively (Table S3). The hygromycin resistance gene (*E. coli hphB*) was amplified with primers OAM597/OAM598. These three amplicons were then recombined by double-joint PCR as described [47]. The extension product was amplified by PCR with the nested primers OAM1159/OAM1160 which contain *Bam*HI restriction sites added at the ends. This product was cloned into pGEM®-T Easy (Promega) and the insert in plasmid AM1538 was sequenced to verify the construct. The gene replacement cassette was cloned as a *Bam*HI fragment into AM1145, the binary vector for *Agrobacterium tumefaciens* transformation [48], creating AM1563. AM1563 was transformed into *A. tumefaciens* strain AD965. *C. posadasii* strain Silveira was transformed with AD965/AM1563 as previously described [48] and hygromycin resistant strains selected and passaged three times to get homokaryons, prior to molecular analysis. Gene replacement strains and ectopic insertion transformants were identified by Southern hybridization analysis as described previously [48], or by PCR. For PCR analysis, primers OAM597 and OAM598 were used to confirm the presence of the hphB gene in all transformants. Primers OAM1611 located upstream of the *RYP1* locus and OAM431 of the *hphB* gene were used to detect a homologous integration event of the transforming construct at the 5’ end of *RYP1*, and primers OAM1612 located downstream of the *RYP1* locus along with OAM432 were used to detect homologous integration of the transforming construct at the 3’ end of *RYP1*. For initial experiments including *in vitro* growth, Δ*ryp1* strains 1563.1, 1563.4, 1563.7 and 1563.14 were used along with ectopic transformed line 1563.19, where the *RYP1* deletion construct was integrated elsewhere in the genome. For mouse virulence and protection studies, Δ*ryp1* strain 1563.7 was used. For further experiments to validate mouse virulence results, an additional set of strains were used; Δ*ryp1* strains 1563.7, 1563.10, 1563.14 and 1563.15 and ectopic transformed strain 1563.1.

### RNA-seq library preparation and sequencing

For expression studies, total RNAs were isolated from WT and Δ*ryp1* hyphal and spherule cultures. Duplicate hyphal cultures of each strain were grown by inoculation of 5 x 10^7^ arthroconidia into 100 ml of 2X GYE and then incubated with shaking at 180 rpm for 48 h at 28°C. Spherule cultures of WT and Δ*ryp1* were also grown in duplicate by inoculating 100 ml of modified Converse liquid medium with 3 x 10^8^ arthroconidia and growing them at 38°C with 20% CO_2_, with shaking at 180 rpm for 48 h [49]. RNAs were isolated using a modified acid-phenol preparation as described previously [49] with the modification that liquid N_2_ grinding was used to initially disrupt the cells. RNAs were resuspended in diethyl pyrocarbonate-treated dH_2_O prior to assessment of their quality and concentration with an Agilent Bioanalyzer (Agilent Technologies, Palo Alto, CA).

RNA-seq libraries were prepared as previously described [25]. Briefly, for the isolation of mRNAs, five μg total RNA from each sample was subjected to poly-A purification using Dynabeads Oligo (dT)_25_(Invitrogen-ThermoFisher). RNA libraries were prepared using NEBNext Ultra Directional RNA Library Prep Kit (NEB). Quality of the total RNA, mRNA and library was confirmed using Bioanalyzer (Agilent). Sequencing was done in-house at UCSF Center for Advanced Technology using Illumina HiSeq2500 instruments. 11 to 19 million single-end 50 bp reads were obtained for each sample.

### RNA-seq data analysis

Predicted mRNA sequences for *Coccidioides posadasii* Silveira were downloaded from the Broad Institute on 3/18/2015 (http://www.broadinstitute.org/annotation/genome/coccidioides_group/MultiDownloads.html)

[23,50].

Relative abundances (reported as TPM values [51]) and estimated counts (est_counts) of each transcript in each sample were estimated by alignment free comparison of k-mers between the preprocessed reads and mRNA sequences using KALLISTO version 0.46.0 [52]. Further analysis was restricted to transcripts with TPM ≥ 10 in at least one sample.

Differentially expressed genes were identified by comparing replicate means for contrasts of interest using LIMMA version 3.30.8 [53,54]. Genes were considered significantly differentially expressed if they were statistically significant (at 5% FDR) with an effect size of at least 2x (absolute log2 fold change ≥ 1) for a given contrast.

### Murine virulence and protection studies

To assess the pathogenicity of Δ*ryp1* mutants, 8-week old female C57BL/6 mice (B6) were anesthetized with ketamine/xylazine and infected intranasally (IN) as previously described [55]. Briefly, 8-10 mice per group were challenged with 50 or 1000 spores of the Δ*ryp1* mutant strains. As a control, B6 mice were given 50 WT spores, and in some studies, B6 mice were infected with 50 spores of a transformed line with an ectopic insertion of the *RYP1* gene deletion construct.

The potential for Δ*ryp1* strains to protect mice from subsequent infection by WT was assessed by performing vaccination experiments as described for Δ*cps1* [19]. Six-week old B6 mice were primed with 2.5 x 10^4^ Δ*ryp1* (strain 1563.7) arthroconidia either by intraperitoneal (IP) or subcutaneous (SC) injection of spores in groups of eight mice and two weeks later boosted with the same number of spores. Two other groups of mice were used as a positive control for vaccination by being vaccinated in a similar manner with Δ*cps1* arthroconidia, at a dose of 5 x 10^4^ spores. As a control, mice were vaccinated with a *S. cerevisiae* culture supernatant, which was previously used as an adjuvant for a chimeric protein antigen that shows some vaccine protection for *Coccidioides* [56]. Mice were challenged four weeks later with 90 WT arthroconidia and sacrificed two weeks later for lung fungal burden determination. Spleens were cultured whole to determine dissemination [56]. All murine infections and handling were carried out at animal BSL3 facility and all procedures with mice were approved by the University of Arizona Institutional Animal Care and Use Committee. Animals were housed and cared for according to PHS standards.

### Other software and libraries

We wrote custom scripts and generated plots in Prism (GraphPad Software, San Diego, CA), R 3.3.3 [57] and Python 2.7.13, using Numpy 1.12.1 [58] and Matplotlib 2.0.0 [59]. Jupyter 4.2.3 [60] notebooks and JavaTreeView 1.1.6r4 [61] were used for interactive data exploration.

## Data Availability

All relevant data are contained within the paper and/or Supporting Information files. For high-throughput sequencing data, the raw data are available at the NCBI Gene Expression Omnibus (GEO) databases under GEO accession GSE178277.

## Supporting Data

**S1 Table. Table of Kallisto quantification, limma statistics, and annotations for differentially expressed genes**. Excel-compatible tab-delimited text conforming to JavaTreeView extended CDT format. Each row is a transcript with the UNIQID column giving the Broad Cp Silveira systematic gene name. The NAME column gives manually curated short gene names, transferred from *Histoplasma capsulatum* G217B (HcG217B, GenBank GCA_017607445.1) augmented with *Coccidioides*-specific gene names. The next three columns give limma BH-adjusted p-values for differential expression in each of the three contrasts. The next 8 columns give kallisto estimated counts for each sample. The next 8 columns give kallisto normalized abundances (TPMs) for each sample. Cp_GenBank and Cp_anno give the GenBank Cp Silveira accession and annotation respectively. CiRS, CiRS_GenBank, and Ci_anno give the Broad systematic gene name, GenBank accession, and annotation respectively for the InParanoid-mapped *C. immitis* RS ortholog. HcG217B, HcG217B_anno, and HcG217B_GenBank give the GSC systematic gene name and annotation (from Voorhies et al, submitted) and GenBank accession for the InParanoid-mapped HcG217B ortholog. Cp_new_GenBank gives the GenBank accession for the corresponding gene in the newly available Cp Silveira assembly and annotation (BioProject PRJNA664774) [62]. Class indicates the differential expression pattern, as referenced in the results and discussion sections. BGCOLOR gives the hex code for the class coloring in Fig 3. GWEIGHT is a place-holder column for JavaTreeView compatibility. The final three columns give the limma fit values for the three contrasts plotted in Fig 3.

**S2 Table. Table of Kallisto quantification, limma statistics, and annotations for all expressed genes**. Excel-compatible tab-delimited text conforming to JavaTreeView extended CDT format. Columns are exactly as for S1 Table, but rows include all 9711 analyzed genes. The estimated counts in this file are sufficient to recapitulate the limma analysis.

**S3 Table. Table of oligonucleotides used for Δ*ryp1* mutant creation and analysis**.

## Acknowledgements

We thank members of the Beyhan, Galgiani, Orbach, and Sil labs for helpful discussions.

